# M2 receptors are required for spatiotemporal sequence learning in mouse primary visual cortex

**DOI:** 10.1101/2022.02.09.479792

**Authors:** Susrita Sarkar, Catalina Martinez Reyes, Cambria M. Jensen, Jeffrey P. Gavornik

**Author notes:** The authors declare no competing financial interests.

## Abstract

Acetylcholine is a neurotransmitter that plays a variety of roles in the central nervous system. It was previously shown that blocking muscarinic receptors with a non-selective antagonist prevents a form of experience-dependent plasticity termed “spatiotemporal sequence learning” in the mouse primary visual cortex (V1). Muscarinic signaling is a complex process involving the combined activities of five different G-protein coupled receptors, M1-M5, all of which are expressed in the murine brain but differ from each other functionally and in anatomical localization. Here we present electrophysiological evidence that M2, but not M1, receptors are required for spatiotemporal sequence learning in mouse V1. We show in male mice that M2 is highly expressed in the neuropil in V1, especially in thalamorecipient layer 4, and co-localizes with the soma in a subset of somatostatin expressing neurons in deep layers. We also show that expression of M2 receptors is higher in the monocular region of V1 than it is in the binocular region, but that the amount of experience-dependent sequence potentiation is similar in both regions, and that blocking muscarinic signaling after visual stimulation does not prevent plasticity. This work establishes a new functional role for M2-type receptors in processing temporal information and demonstrates that monocular circuits are modified by experience in a manner similar to binocular circuits.

**Significance Statement:** Muscarinic acetylcholine receptors are required for multiple forms of plasticity in the brain and support perceptual functions, but the precise role of the five subtypes (M1-M5) are unclear. Here we show that the M2 receptor is specifically required to encode experience-dependent representations of spatiotemporal relationships in both monocular and binocular regions of mouse V1. Muscarinic receptors are metabotropic in nature and have complex downstream signaling effects that can be difficult to isolate experimentally. In addition to identifying a novel functional role for M2 receptors, this work also suggests that V1 circuits can be used as an in-vivo model to understand the relationship between cholinergic signaling and coding of complex temporal relationships in cortical circuits.

## Introduction

Acetylcholine (ACh) is involved in a wide range of brain functions including attention (Herrero et al., 2008; Parikh et al., 2007; Sarter et al., 2005), memory formation (Hasselmo, 2006), sensory perception (Pinto et al., 2013; Goard & Dan, 2009), temporal processing (Liu et al., 2015), plasticity (Bear and Singer, 1986), and oscillatory synchronization (Gu, 2003; Hasselmo and McGaughy, 2004; Steriade, 2004; Thiele, 2013). The complex relationship between cholinergic signaling and neural activity is attributed to functional differences and anatomical distributions of nicotinic (nAChR) and muscarinic (mAChR) receptors. Of interest here are the mAChRs which are required for a form of experience-dependent plasticity that can encode spatiotemporal information in mouse V1 (Gavornik and Bear, 2014) and other areas as well (Sidorov et al., 2020; Finnie et al., 2021).

The 5 subtypes of mAChRs (M1-M5) are functionally classified into M1-type (M1, M3 and M5, coupled to G_q/11_) and M2-type (M2 and M4, coupled to G_i/o_) receptors (Thiele, 2013; Groleau et al., 2015; Coppola and Disney, 2018). All but M5 are expressed to varying levels in the cortex, striatum, thalamus, and hippocampus (Bubser et al., 2012.; Levey et al., 1991; Thiele, 2013; Volpicelli & Levey, 2004). M1-type receptors are predominantly excitatory and post-synaptic whereas M2-type receptors are generally presynaptic (Thiele, 2013). Pre-synaptic M2-type receptors can prevent neurotransmitter release at cholinergic (Mrzljak et al., 1993; Groleau et al., 2015) and GABAergic (Salgado et al., 2007) terminals. Individual studies report information about the cell-type and layer-specific localization of muscarinic receptors in V1, though not all those studies were conducted in mice.

In V1 of primates such as the macaque monkey, a large extent of the cholinergic modulation is aimed at GABAergic neurons such as parvalbumin (PV) neurons, 87% of which express M1 receptors and 31% of which express M2 receptors (Disney and Aoki, 2008). M2 receptors represent 36% of the total mAChRs in primate V1 (Flynn et al., 1995). In V1 (area 17) of the rhesus monkey, M1 is highly concentrated in superficial layers and less so in the middle cortical layers but M2 is highly concentrated in layer IVC of the rhesus monkey and superficial layers of the macaque monkey (Lidow et al., 1989; Tigges et al., 1997; Gu, 2003). M2 receptors are located at presynaptic sites in primate V1 where they function as heteroreceptors and modulate excitatory synaptic transmission (Mrzljak et al., 1996). M1 receptors are found at postsynaptic sites such as spines and dendrites in primates, where they regulate both cholinergic and excitatory synaptic transmission (Mrzljak et al., 1993; Volpicelli and Levy, 2004). M1 and M2 receptors are the two most abundant ones in the rodent visual cortex whereas M1, M2 and M4 receptors are predominant in human and primate V1 (Flynn et al., 1995; Groleau, et al., 2015). In humans and rodents, M1 receptors are generally found at postsynaptic sites in pyramidal cells such as the soma and dendrites but in primates they are found to a great extent on GABAergic neurons (Mrzljak et al., 1993; Gu, 2003; Gulledge et al., 2009; Groleau et al., 2015; Disney et al., 2006). In rhesus macaque monkeys, attentional modulation in V1 is brought about by muscarinic receptors and is blocked by scopolamine, the non-selective muscarinic antagonist (Herrero et al., 2008). In primate V1, ACh is believed to modulate lateral connections between neurons with similar tuning properties (Ramalingam et al., 2013; Herrero and Thiele, 2021).

Scopolamine is a non-selective antagonist that blocks M1-M5 (Lochner and Thompson, 2016). In a 2014 study, Gavornik and Bear showed that binocular V1 can encode the spatiotemporal content of visual sequences, but that this form of plasticity is prevented by scopolamine. This work is not the only to show that ACh can entrain temporal representations in V1. Cholinergic signaling also allow circuits in rodent V1 to learn temporal intervals in both in vivo (Shuler & Bear, 2006; Liu et al., 2015; Namboodiri et al., 2015) and ex vivo preparations (Chubykin et al., 2013). It remains an open question which specific receptor populations are required for this plasticity (Muñoz & Rudy, 2014). Here we employ the same “sequence learning” paradigm used in Gavornik and Bear, 2014, which causes responses evoked in V1 by a 4-element visual sequence to potentiate with exposure and is somewhat selective for the temporal order of sequence elements. We find that spatiotemporal sequence learning occurs normally when M1 receptors are blocked and is prevented by antagonizing M2 receptors. We also report on the layer-specific distribution of muscarinic receptors in V1. Most M2 expression is found in the neuropil, but it is also expressed somatically on some somatostatin (SST) positive neurons in deep cortical layers, and M2 expression is higher in monocular V1 than in the binocular region. Despite this, we show that sequence learning occurs with similar potentiation magnitude in monocular V1 as it does in binocular V1. Finally, we show that blocking receptors immediately after visual stimulation does not prevent plasticity.

## Methods and Materials

### Animals

Male C57BL/6 mice (Charles River Laboratories) were group housed with littermates (four mice per cage) on a 12-h light/dark cycle and provided food and water *ad libitum*. Experimental and control groups were selected randomly from littermates and yoked throughout experiments. All experiments were performed during the light-cycle and animals were used for a single experiment only. All procedures were approved by the Institutional Animal Care and Use Committee of Boston University.

### Electrode implantation

Mice were anesthetized with an intraperitoneal (IP) injection of 50 mg per kg ketamine and 10 mg per kg xylazine and prepared for chronic recording. To facilitate head restraint, a steel headpost was affixed to the skull anterior to bregma using cyanoacrylate glue. Small (<0.5 mm) burr holes were drilled over binocular visual cortex (3 mm lateral from lambda) or monocular visual cortex (::2.2 mm lateral from lambda) and tungsten microelectrodes (for field recordings, FHC) were placed 450 μm below the cortical surface. In all surgeries, a reference electrode (silver wire, A-M systems) was placed in the dura over parietal cortex. All electrodes were rigidly secured to the skull using cyanoacrylate glue. Dental cement was used to enclose exposed skull and electrodes in a protective head cap (Metabond and Ortho-jet). Buprenex (0.1 mg per kg) was injected subcutaneously for postoperative pain amelioration. Surgery was performed around postnatal day 45. Mice were monitored for signs of infection and allowed at least 24 h of recovery before habituation to the recording and restraint apparatus and were excluded from experiments in the event of unsuccessful electrode implantation (lack of visual response, incorrect depth, poor grounding, etc.).

### Stimulus presentation

Visual stimuli were generated using custom software written in Matlab (MathWorks) using the PsychToolbox extension (http://psychtoolbox.org/) to control stimulus drawing and timing. Sequences were constructed of four elements and an inter-sequence gray period. Each element consisted of a full-screen oriented high-contrast sinusoidal grating (0.5 cycles per deg). Sequence elements were separated by a minimum of 30 degrees and the order was restricted to prevent the appearance of rotation. Each element was held on the screen for 150 ms and sequences were separated by 1.5 s of gray screen. Grating stimuli spanned the full range of monitor display values between black and white, with gamma correction to ensure a linear gradient and constant total luminance in both gray-screen and patterned stimulus conditions. During experiments, animal handling involved placing each mouse (regardless of group membership, of which the investigator was aware) into the head-fixed presentation apparatus. Each sequence was presented 200 times per day in four groups of 50 presentations with each group separated by 30 s. In the monocular versus binocular sequence learning experiment, monocular cortex was isolated by offsetting the screen to the right and adjusting it to an approximately 45° angle relative to the mouse’s midline while also covering the left eye with an opaque occluder for the first 5 days. Isolation was confirmed by showing brief flashing visual stimuli and verifying evoked responses were evident only in the in left monocular cortex (with the screen directly in front of the mouse, ipsilateral projections from the right eye drive responses in the right binocular cortex). For binocular training on days 6-10 the screen was placed directly in front of the mice and both eyes were open.

### Data recording, analysis, and presentation

All data was amplified and digitized using the digital Recorder-64 system (Plexon). Local field potentials (LFPs) were recorded with 1-kHz sampling and a 200-Hz low-pass filter. LFP voltage traces in all figures show the average response of all mice in an experimental cohort. Data was extracted from the binary storage files and analyzed using custom software written in C++ and Matlab, all of which is available for download at gavorniklab.bu.edu/supplemental-materials. To quantify plasticity effects across days, sequence magnitude was defined as the average response magnitude (algorithmically scored trough-to-peak within the first 150 ms of visual stimulus onset) for each of the four elements in a sequence. Ordinal effects were assessed using responses to the 2nd and 3rd elements (Finnie et al., 2021).

### Scopolamine injections

For the post-stimulus scopolamine experiment, mice were intraperitoneally injected with either 3 mg per kg of scopolamine hydrobromide (Tocris) dissolved in saline or vehicle (saline). Injections were performed immediately after sequence presentation on experimental days 1–4. This procedure was chosen to match previously published reports demonstrating that this dose is sufficient to effectively block muscarinic receptors in vivo (Gavornik & Bear, 2014). To allow sufficient time for drug washout, recordings on the 5th day occurred at least 24 h after the last injection on day 4.

### VU 0255035 injections

For the systemic VU 0255035 experiment, experimental mice were intraperitoneally injected with 10 or 25 mg per kg of VU 0255035 (R&D Systems) in DMSO + saline. Vehicle control animals were injected with DMSO + saline. Injections were performed 30 min before sequence presentation on experimental days 1–4. This procedure was chosen to match previously published reports demonstrating that this dose is sufficient to effectively block M1 receptors in vivo (Sheffler et al, 2009; Crans et. al, 2020). To allow sufficient time for drug washout, recordings on the 5th day occurred at least 24 h after the last injection on day 4.

### Open-Field Exploration

To quantify the effect of drug on locomotion, mice were placed in a 40×40×30cm box (Maze Engineers) and allowed to freely explore. Video was acquired from above using a Microsoft Kinect camera and analyzed in real-time track mouse movements (all code available at gavorniklab.bu.edu). Mouse location and height data were stored in a text file. To determine the effect of the M1 antagonist VU 0255035, we analyzed stored position data to calculate total distance traveled during 30-minute sessions following IP injections of vehicle or 10 mg/kg, 25 mg/kg, and 50 mg/kg of the drug. Each mouse was treated with a single concentration on any day, and each mouse was injected with each concentration over the course of the experiment (injection doses were randomized by day).

### AQ-RA 741 infusions

For the local AQ-RA 741 experiment, a 26-gauge guide cannula (P1 Technologies) was implanted lateral to the recording electrode in one hemisphere, angled at 45° relative to the electrode and positioned slightly below the cortical surface. Guide cannulae were affixed to the skull with cyanoacrylate glue and encased in the dental cement head cap. Dummy cannulae were installed to protect against external debris entering the guide cannulae. After several days of recovery and habituation to the head-fixed restraints, the dummy cannulae were removed and infusion cannulae were lowered into the guides (see schematic in Fig. 2A). A Nanoject II (Warner Instruments) was used to infuse 1 μl of vehicle (DMSO + saline) or AQ-RA 741 (R&D SYSTEMS, in DMSO + saline) before stimulus presentation on experimental days 1 and 2 under isoflurane anesthesia. Animals were recorded 30 minutes after infusion completion to ensure full recovery from the anesthesia. Mice received either drug or vehicle infusions, and all mice that evidenced clear visual responsiveness (that is, no trauma caused by infusion) were included in the study. Local treatment was performed for only 2 days to minimize inherent cortical trauma associated with repeated insertions and removals of the infusion and dummy cannulae; training continued for 2 additional days after drug washout to verify that V1 was rendered aplastic because of drug treatment and not an unintentional cortical lesion.

### Immunofluorescence

Animals were intraperitoneally injected with pentobarbital (100 mg/kg, Fatal Plus, Covetrus, 035946) and then perfused with 4% paraformaldehyde in PBS. Fixed tissue was cryoprotected in 30% sucrose in water and sectioned via cryostat to obtain coronal sections (50*β*m slices). Sections were permeabilized and blocked using a solution of 0.1% Triton-X, 10% NGS, 1% BSA in PBS. Sections were incubated with primary antibodies against M1 muscarinic receptor (rabbit anti-M1, Novus Biologicals, NBP1-87466), M2 muscarinic receptor (rat anti-M2, Millipore, MAB367), Parvalbumin (PV) (rabbit anti-PV, Abcam, ab11427) or Somatostatin (SST) (rabbit anti-SST, Fisher Scientific, NBP1-87022) neuronal proteins at a dilution of 1:500 each in 10% NGS in PBS for 48 hours at 4°C except the M1 antibody for which we used a 1:50 dilution. Slices were next washed three times with 0.1% Triton-X in PBS for 5-10 minutes per wash, and then incubated with secondary antibodies at 1:500 dilution in either 10% NGS in PBS (goat anti-rat, Alexa Fluor 647, Thermo Fisher, A21247) or 1% BSA in PBS (donkey anti-rabbit, Alexa Fluor 488, Thermo Fisher, A21206) at room temperature for 2 hours. For nuclear dsDNA staining, slices were incubated in a 1:10,000 dilution of Hoechst (Thermo Fisher, H1399) in 1% BSA in PBS for 15 minutes at room temperature. Following secondary antibody incubation, slices were rinsed three times in PBS for 5-10 minutes per rinse. Slices were then mounted to slides with Fluoro-gel mounting media. Fluorescence images were captured using a Nikon Eclipse Brightfield Microscope in the following channels: DAPI (Hoechst/blue), FITC (Alexa 488/green), CY5 (Alexa 647/red).

To quantify the number of PV+ and SST+ neurons, we analyzed histological images from 50 µm slices of a single hemisphere of V1 (including monocular and binocular regions) in ImageJ using the following steps: 1. grayscale conversion 2. background subtraction 3. threshold adjustment to remove noise 4. conversion to black and white format 5. cell counting. For PV/SST and M2 co-localization, we first counted the number of interneurons (green channel) and M2 positive cells (red channel) separately, then counted the number of interneurons that showed M2 expression by looking for yellow cells (red-green overlap) in a composite image.

To quantify M1 and M2 expression levels, histological images were imported into MATLAB and converted to grayscale. The average pixel intensity of V1 was computed. Each image was then cropped into monocular and binocular V1 (position determined using hippocampal landmarks identified in the Allen Mouse Brain Atlas) and processed independently. The row-wise mean of each monocular and binocular image was calculated, creating an array of values corresponding to the average pixel intensity by cortical depth. These values were normalized by the average pixel intensity of the corresponding V1 image. The normalized pixel intensity values were averaged among different animals and plotted against cortical depth (values extracted from metadata in ImageJ) along with 95% confidence intervals of the group mean. To compare expression levels in supragranular, granular, and infragranular layers, mean intensity levels were calculated for 0-350, 350-450, and 450-700 µm ranges.

### Experimental Design and Statistical Analyses

All experiments were designed following the protocols established in Gavornik and Bear, 2014 and used elsewhere (Sidorov et al., 2020; Finnie et al., 2021). The primary goal was to test the relationship between drug treatment and sequence potentiation, which we did by exposing drug-treated experimental groups with vehicle-treated controls and multiple time points during and after training. One experiment (Fig. 5) was designed to compare potentiation occurring in the monocular and binocular cortex and did not involve drug treatment. All statistics were performed using the SPSS software package. Unless otherwise noted, comparisons between groups were made using a mixed model repeated measures ANOVA (RM-ANOVA), where session (e.g., recording day) was compared within subject groups assuming sphericity (confirmed with Mauchly’s test) and treatment was a between-subject variable. When main effects or interaction terms were significant, pairwise comparisons were performed in SPSS with the Bonferroni correction for multiple comparisons. When making direct comparisons of mean response levels between groups, as when comparing ABCD to DCBA, two-tailed paired-sample t-tests were used to calculate significance. The size of experimental cohorts was planned based on previously published experiments and our own experience which indicates that a minimum of 5 animals is needed in each cohort to reach statistical significance with adequate power, and we used power estimates in SPSS to verify that all statistically significant effects were ≥ 0.8 for α=0.05. In all cases, we planned the experiments using the minimum number of mice expected to be required to achieve significant results based on our expectation of attrition rates and effect sizes (more mice were used for the M2 experiments because we anticipated that the cannulation technique would lead to a higher rate of surgical failures than it did).

## Results

### M1 receptors are not required for spatiotemporal sequence learning in mouse V1

Of the M1-type receptors, only M1 is abundant in rodent V1; previous studies indicate that M3 is not detectable in rodent V1 via immunocytochemistry (Groleau et al., 2015; Levey et al., 1994) and M5 is found on endothelial cells with minimal expression in rodent visual cortex (Elhusseiny and Hamel, 2000; Groleau et al., 2015). Both M2-type receptors are found in the mouse brain. M2 is dominate in the cortex and is found in rodent V1, while M4 dominates in the striatum and has not been reported in V1 (Groleau et al., 2015; Bubser et al., 2012; Levey et al., 1991; Thiele, 2013; Zhang et al., 2002). M1, which is generally excitatory, seemed more likely than M2 to be responsible for potentiated responses associated with spatiotemporal plasticity. Accordingly, for our first experiment we pharmacologically blocked M1 receptors while using chronically-implanted electrodes implanted in binocular layer 4 of mouse V1 to record visually evoked potentials (VEPs) in response to a sequence stimulus. Mice were randomly assigned to experimental or control groups and trained for 4 days via passive exposure to the sequence ABCD (see methods, Fig. 1A) On each training day, mice were injected intraperitoneally (IP) with either vehicle (control) or a highly selective M1 antagonist VU 0255035 (experimental, 10 mg/kg) 30 mins before stimulus presentation. The VU 0255035 dose used was chosen to mirror previous reports that this concentration is sufficient to block receptors *in vivo* (Sheffler et al., 2009; Crans et al., 2020; Weaver et al., 2009) and prevent some forms of visual learning (Rahman et al., 2020). On the test day (day 5), both groups were presented with the same stimulus without a prior injection. VEP magnitudes increased with training in both groups and were markedly larger on day 5 than on day 1 (Fig. 1B) with no obvious difference between groups on any recording day. There is no significant difference in sequence magnitudes (quantified as the average trough-to-peak response of each element, see methods) as a function of treatment group (Fig. 1B-E).

**Figure 1.**
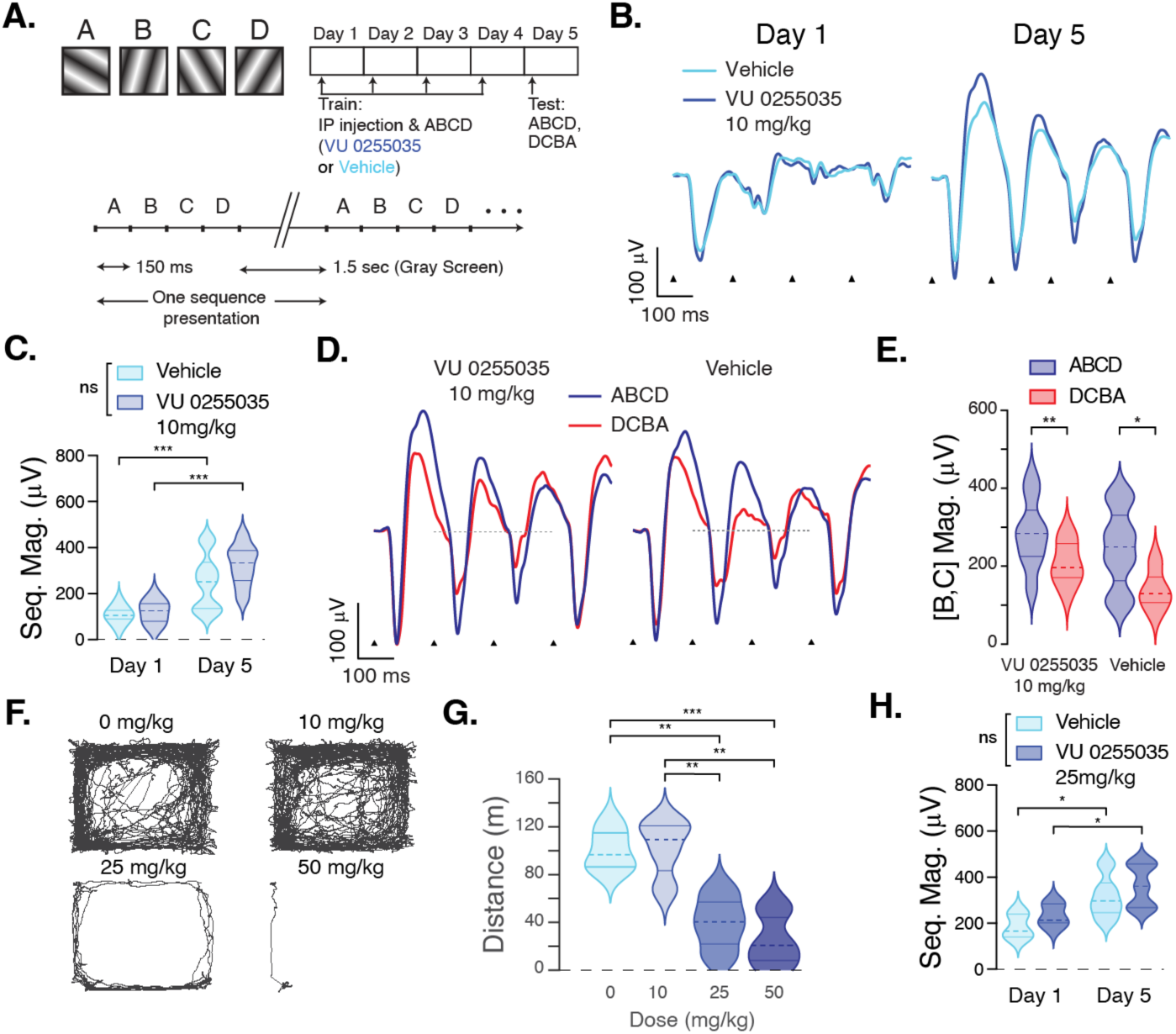
M1 receptors are not required for spatiotemporal sequence learning in V1. **A.** Mice were treated with either the M1 receptor antagonist VU 0255035 (exp, 10 mg/kg, *n* = 8) or vehicle (ctrl, *n* = 5) on each of four training days. Training consisted of 200 presentations of the sequence ABCD. All mice were tested using ABCD and reordered sequence DCBA on day 5. **B.** Sequence-evoked LFPs looked very similar in drug and vehicle-treated animals at baseline (day 1) and training-induced potentiation (day 5). Voltage traces represent stimulus locked LFP response averaged across all mice in each group, triangles mark the onset of each stimulus in the sequence. **C.** Sequence magnitudes (quantified as the average trough-to-peak response to each element) recorded on days 1 and 5 were analyzed using a mixed model RM-ANOVA which showed a significant effect of recording day (*F*_1,11_ = 71.659, *p* < 0.001) but no effect of treatment (*F*_1,11_ = 1.071, *p* = 0.323) or significant interaction between treatment and day (*F*_1,11_ = 2.272, *p* = 0.160). Post-hoc analysis with a Bonferroni correction revealed that sequence magnitude was significantly larger on day 5 than day 1 in both drug (202.1 µV, p < 0.001) and vehicle (141.0 µV, *p* = 0.001) treated animals. Quantified data is visualized using violin plots whose shapes represent an estimate of the probability density distribution (solid horizontal lines mark the first and third quartile divisions, the dashed line shows the median). **D.** Average responses to ABCD (blue) are larger than those to DCBA (red) in drug (left) and vehicle (right) treated mice. **E.** To avoid the confoundingly large response that always occurs following the transition from a gray screen to patterned stimuli, we chose to assess ordinal effects using only responses to the 2nd and 3rd elements (BC or CB, marked with gray dashed line in D, results are similar when calculated from the entire sequence). A 2-way ANOVA shows that there is a significant effect of order (*F*_1,20_ = 7.121, *p* = 0.015) but not drug treatment (*F*_1,20_ = 2.395, *p* = 0.137) and no significant interaction between the two (*F*_1,20_ = 0.206, *p* = 0.655). Post-hoc comparisons show that ABCD is significantly larger than DCBA in both drug (76.0 µV, t(7) = 4.958, *p* = 0.002) and vehicle (87.8 µV, t(4) = 2.768, *p* = 0.050) treatment groups. To verify drug action, mouse (*n =* 4) position was tracked during a 30-minute period of free exploration following I.P. injections of vehicle, 10, 20, or 50 mg/kg VU 0255035. **F.** Representative plot of a single mouse’s position following injections demonstrates locomotion decreases with drug dose. **G.** Total distance traveled is significantly higher with vehicle or 10 mg/kg relative to 25 or 50 mg/kg (Within subject effective of dose *F*_3,6_=10.104, *p* = 0.009. Bonferroni corrected t-tests: 0vs25 t(6)=4.167 *p*=0.001, 0vs50 t(5)=4.773 *p*<0.001, 10vs25 t(6)=3.634 *p*=0.002, 10vs50 t(5)=4.028 *p*=0.002. Other comparisons were not significant at 0.05) **H.** The experiment from panel A was repeated with 25 mg/kg VU 0255035 (*n =* 5 for both treatment groups). As with lower dose, there was a significant effect of recording day (*F*_1,8_ = 29.037, *p* < 0.001) but no effect of treatment (*F*_1,8_ = 1.310, *p* = 0.285) or significant interaction between treatment and day (*F*_1,8_ = 0.000, *p* = 0.990).

This negative result could potentially be explained by experimental error (drug batch issues, dilution errors, etc.) rather than M1 receptor blockade. To verify drug action, we designed an experiment to quantify our observation that animals injected with VU 0255035 show reduced motility by tracking mouse location in an open arena following drug injection using an overhead camera (see methods). Drug concentration affected locomotion in a dose dependent manner, with significant declines in total distance traveled over a 30-minute time periods relative to vehicle at both 25 and 50 mg/kg (Fig. 1F-G). Since there was no effect at 10 mg/kg, we repeated the sequence learning experiment using 25 mg/kg. Even at this dose, 2.5 times higher than the dose shown by Rahman, et al. to block memory formation *in vivo*, there was no difference between drug and vehicle treated animals (Fig. 1H), confirming that M1 receptors are not required for sequence potentiation in V1.

### M2 receptors are required for spatiotemporal sequence learning in mouse V1

Our next experiment was to block M2 receptors using a high affinity, selective M2 antagonist AQ-RA 741 (Doods et al., 1991; Dorje et al., 1991). AQ-RA 741 has higher affinity for the M2 and M4 receptors compared to other muscarinic receptors (Watson & Eglen, 1999; Dorje et al., 1991) making it an important tool to differentiate the action the M2-type receptors. Unlike VU 0255035, AQ-RA 741 is unable to cross the blood brain barrier (Doods et al., 1993) under physiological conditions. Accordingly, we modified our experimental design to infuse the drug directly into V1 via indwelling canulae (Fig. 2A) while recording sequence VEPs from binocular layer 4 of mouse V1. As before, mice were randomly assigned to experimental and yoked control groups. On each training day, mice were intracerebrally infused with AQ-RA 741 or vehicle under light isoflurane anesthesia and exposed to stimulation 30 mins later. As shown in Fig. 2B,D, there was minimal change in experimental VEPs over this period while VEPs in control animals potentiated considerably. To ensure that the lack of potentiation in experimental animals was caused by drug action and not damage to cortical tissue, we continued to train the experimental cohort for an additional two days. As shown in Fig. 2C-D, VEPs in experimental animals on day 3 (the first day of non-drug treated visual exposure) were essentially the same as on day 1 in control mice and showed normal sequence potentiation to ABCD on days 4 and 5. While VEP magnitude changes were significantly different between experiment and control animals on days 2 and 3 during drug treatment, there was no significant difference when comparing the groups on the 3rd day of non-treated potentiation. The response to the learned sequence was also larger than a novel sequence in both groups (Fig. 2E-F). We interpret these results to mean that significant potentiation between days 1 and 3 in drug-treated animals was prevented by drug action. Once drug administration was stopped after day 2, treated tissue regained the capacity to potentiate to ABCD.

**Figure 2.**
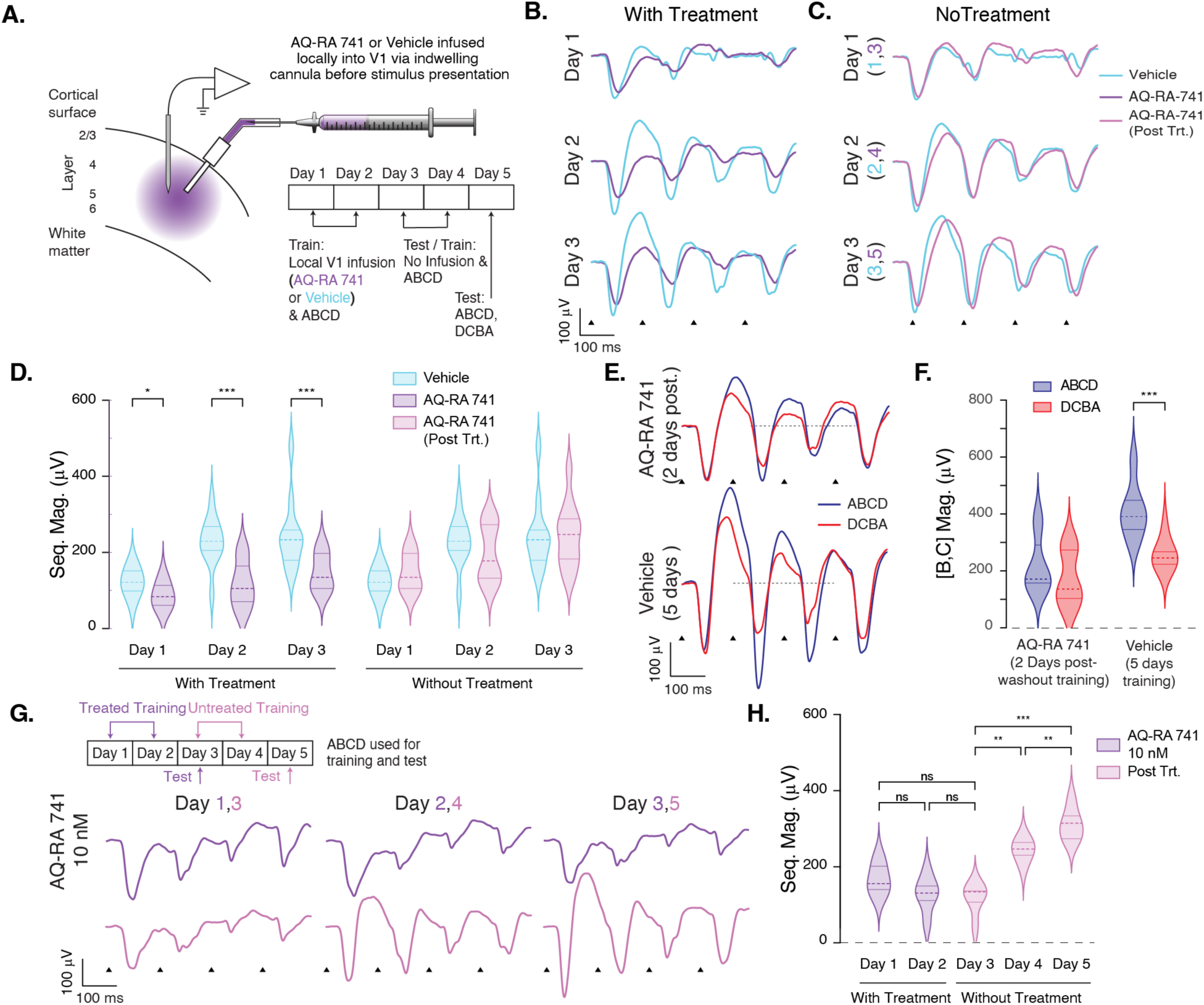
M2 receptors are required for spatiotemporal sequence learning in V1. **A.** Mice were treated with intracerebral infusions of the M2 antagonist AQ-RA 741 (∼ 400 μM) via an indwelling cannula surgically implanted in V1. Animals were treated with either drug (exp, *n* = 12) or vehicle (ctrl, *n* = 11) before stimulus presentation on training days 1 and 2 but not before test stimuli on day 3. Training continued for an additional day and a final set of test stimuli (ABCD for all animals, ABCD and DCBA in 7 vehicle and 8 drug treated) was presented on day 5. **B.** Sequence-evoked responses potentiated normally in vehicle-treated mice but were impaired in drug-treated animals. **C.** After drug washout, the responses in drug-treated animals potentiated at a level commensurate with vehicle-treated animals. This panel compares responses over the first three days of untreated training for vehicle and drug treated mice (days 1-3 and 3-5, respectively). **D.** The effect of training and treatment was analyzed on days 1-3 using a mixed model RM-ANOVA which revealed significant main effects of day (*F*_2,42_ = 33.384, *p* < 0.001) and treatment (*F*_1,21_ = 15.788, *p* = 0.001) and a significant interaction between the two (*F*_2,42_ = 6.200, p = 0.004). Bonferroni-corrected post-hoc tests showed a small but significant difference between groups on day 1 (34.6 µV, p = 0.022) that increased on days 2 (111.9 µV, p < 0.001) and 3 (97.6 µV, p = 0.004). Responses in drug-treated animals increased on both days 2 and 3 but not significantly (1vs2: 29.6 µV, p = 0.144. 2vs3: 30.8µV, p = 0.178) whereas vehicle-treated animals showed a large amount of potentiation on day 2 (1vs2: 106.9µV, p < 0.001) and modest potentiation on day 3 (2vs3: 16.6µV, p = 0.946). The effects of training without treatment were similarly analyzed using the comparisons from panel C. This revealed a significant effect of training (*F*_2,42_ = 33.790, *p* < 0.001) but no effect of treatment group (*F*_1,21_ = 0.009, *p* = 0.924) or significant interaction (*F*_2,42_=2.167, *p* = 0.127), demonstrating that the lack of potentiation on days 2-3 was an acute byproduct of drug treatment. **E.** ABCD evoked a larger response (blue) than DCBA (red) in both treatment groups on day 5. Note that the experimental design meant vehicle and drug-treated animals were tested after 4 and 2 days of training, respectively. **F.** ABCD was significantly larger than DCBA in vehicle treated animals (68.0 µV, t(6) = 5.922, *p* = 0.001). ABCD was also larger on average in drug treated mice, but this difference was not significant (45.0 µV, t(7) = 1.667, *p* = 0.139) though we expect it would be with additional training. **G.** A second infusion experiment was conducted with a lower drug dose. Animals (*n =* 5) were infused with 10 nM AQ-RA 741 before stimulus presentation on training days 1 and 2 and tested without drug on day 3. Training continued for an additional day and animals were again tested on day 5. Sequence-evoked responses did not potentiate with drug present (days 1-3) but potentiated normally following drug washout (days 3-5). **H.** The effect of training and treatment was analyzed using a mixed model RM-ANOVA to compare sequence magnitudes on days 1-3 (potentiation with drug) to days 3-5 (potentiation without drug) which revealed significant effects of both day (*F*_2,16_ = 10.195, *p* = 0.001) and treatment (*F*_1,8_ = 23.278, *p* =0.001) with a significant interaction between the two (*F*_2,16_ = 31.358, *p* < 0.001). Bonferroni-corrected post hoc tests showed that there were no significant changes in VEP magnitude with drug (1vs2: -40.7 µV, *p* = 0.412; 2vs3: -12.0 µV, *p* = 1.000; 1vs3: -52.6 µV, *p* = 0.301) but significant potentiation after washout (3vs4: 128.2 µV, p = 0.002; 4vs5: 68.3 µV, *p* = 0.001; 3vs5: 196.4 µV, *p* < 0.001).

Though AQ-RA 741 has the largest affinity for M2 receptors, it can also bind to M1-type receptors at high enough concentrations. We infused a solution with a molarity of 431 μM. It is difficult to estimate the actual concentration in V1 following dilution in cerebrospinal fluid and molecular diffusion, but it is possible that the drug was binding a substantial fraction of M1 receptors. This would prevent us from concluding that plasticity requires only M2 receptors and not a simultaneous combination of both M1 and M2. To rule out this possibility, we ran a second local infusion experiment using a dilution of 10 nM. This dose was selected to be greater than the inhibitor constant estimated for M2 receptors (Ki ≅ 4.4 nM) while being much lower than the K_i_ estimated for M1 receptors (∼ 62 nM) (Gillberg et al., 1998) and blocked plasticity in a manner like that seen with the higher dose (Fig. 2G-H). Taken together with our finding that M1 is not required for sequence learning, the different functional roles ascribed to M1 and M2 receptors, and the anatomical separation of muscarinic receptors in V1 (see below), these results suggest M2 receptor signaling is necessary and sufficient for spatiotemporal plasticity to occur.

### M2 receptors are differentially expressed based on cortical layer, cell type, and location within V1

Since our electrophysiology experiments revealed that M2 receptors are required for sequence learning in mouse V1, we next turned to immunohistochemistry to look in detail at the layer-wise distribution of M2 receptors in V1. We stained for M2 receptors using immunofluorescent labeling in coronal brain slices acquired from mice age-matched to our experimental cohorts. Resulting images show that M2 receptors are found in all layers of mouse V1 with an intensity profile that varies by layer (Fig 3A). Similar to findings by Ji et al., 2015, expression peaks in layer 4 and is also relatively high in layer 5B (Fig 3B). There is also a marked difference in M2 expression levels between monocular and binocular areas. To confirm that the M2 expression pattern was not attributable to cell density differences, we also stained for dsDNA (DAPI) in the same coronal slices and verified that the expression patterns did not match.

**Figure 3.**
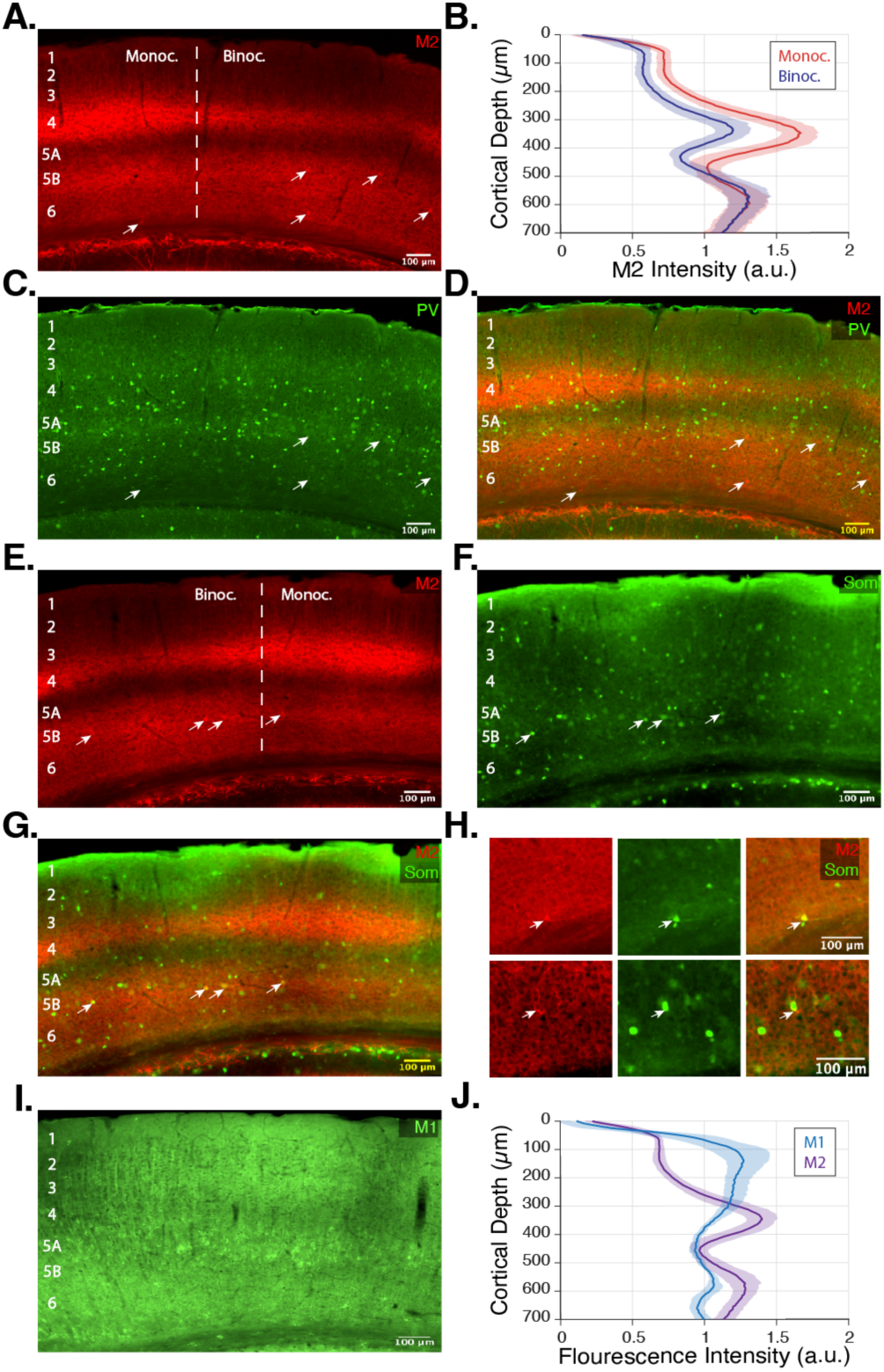
M2 receptors are not expressed on cell bodies of PV interneurons in V1 but are expressed on cell bodies of some SST neurons. **A.** Representative image showing M2 receptor antibody staining. M2 receptors are found throughout V1, though levels vary between layers. Most of the fluorescence originates in the neuropil, though there are a few labeled cell bodies in deep layers (marked with arrows). In addition, there is a clear increase in layer 4 expression in monocular V1 (left) relative to binocular (right, separated by a dashed line, the position of which was determined using hippocampal landmarks identified in the Mouse Brain Atlas). **B.** M2 depth-wise intensity profiles in monocular and binocular V1 quantify higher expression levels in supragranular (17.9%) and granular (45.8%) layers of monocular cortex, while expression in infragranular layers is approximately equal (-1.85%). Solid lines indicate normalized group mean intensity of M2 expression (slices from *n =* 9 mice) with 95% confidence intervals (shaded areas). A 2-way ANOVA of intensity values shows that there is a significant effect of depth (supra/gran/infra, *F*_2,48_ = 40.650, *p* < 0.001) and cortical area (monoc/binoc, *F*_1,48_ = 34.817, *p* < 0.11) with a significant interaction between groups (*F*_2,48_ = 12.316, *p* < 0.001). Post-hoc analysis with Bonferroni correction confirms the previous observation that M2 expression levels are significantly higher in monocular V1 than in binocular V1 in layers 1-3 (0.218 a.u., *p* = 0.001) and layer 4 (0.428 a.u, *p* < 0.001), but the same in deep layers (-0.009 a.u., *p* = 0.884). **C.** Cell bodies identified in the M2 channel (arrows) do not match the location of PV+ somas. **D.** Overlap image of the M2 (red) and PV (green) channel confirms that PV somas do not express M2 receptors. **E.** A second representative image showing M2 expression (details as in A, though from a different mouse and in the left rather than right hemisphere). **F-H.** Anti-SST co-labeling of the same slice reveals that somas located in the M2 channel (red) are also visible in the SST channel (green). We did not find any somas clearly labeled with the M2 signal that did not co-localize with an SST+ neuron. **I.** Representative image showing M1 receptors are densely expressed in the superficial layers 2/3 with somatic labelling in layer 5. **J.** Layer-wise expression pattern of M1 receptors varies relative to M2 receptors as a function of depth (plot shows expression density for monocular and binocular regions combined, slices from *n* = 5 mice).

While most M2 expression was seen in neuropil, we did find a small number of cell bodies that were positive for M2 (arrows, Fig. 3 A, E). Different classes of inhibitory interneurons are functionally implicated in various V1 computations (Chen et al., 2015; Hooks & Chen, 2020; Pfeffer et al., 2013; Niell and Scanziani, 2021) so we decided to determine if M2 receptors are expressed somatically on Parvalbumin (PV) or Somatostatin (SST) interneurons in V1 by using immunofluorescence co-labeling of M2 and PV or SST. We found approximately 116 PV neurons per hemisphere (Fig. 3C, 50-micron slices from n=4 mice, µ=115.5, σ=3.9), none of which co-expressed the M2 label (Fig.3D). We found about 40% fewer SST positive cell bodies in V1 compared to PV (µ=69.8, σ=11.5 SST positive cells per hemisphere, n=4 mice), a small fraction of which do collocate with the M2 signal (4% of SST neurons co-labeled with M2, Fig. 3E-G). The majority, 73%, of M2-positive SST neurons were in the infragranular layers, with the remaining 27% in supragranular layers. All M2-positive somas identified in V1 colocalized with SST stain. The subset of somatically labeled SST neurons might be of particular interest in future studies focused on spatiotemporal plasticity.

Though we did not identify a physiological role for M1 receptors, we wanted to understand their anatomical relationship to M2 receptors. Immunofluorescent labelling (Fig. 3I) shows that M1 receptors are densely expressed in the neuropil of superficial layers 2/3 with robust somatic labelling in layer 5. Plotting the average intensity as a function of depth reveals a clear difference in laminar expression between M1 and M2 receptors (Fig. 3J). Unlike M2 receptors, we find no difference between expression in monocular and binocular areas. This anatomical differentiation between M1 and M2 expression patterns suggests that the receptors play different roles in visual processing and further supports our conclusions from above that M2 receptors are uniquely required for plasticity to occur.

### Experience-dependent sequence potentiation occurs in monocular V1, with potentiation comparable to binocular V1

In the immunofluorescence experiments above, we observed that the level of M2 expression differed between binocular and monocular L4 of mouse V1 (Fig. 3B). All previous sequence learning experiments have been conducted in the binocular region. Reasoning that sequence potentiation might scale with receptor expression density, we hypothesized that monocular V1 would show more plasticity than binocular V1. To test this hypothesis, we implanted electrodes in left monocular cortex (LMC) and right binocular cortex (RBC) and modified our stimulation setup to isolate the LMC to determine whether visual stimulation would drive potentiation similar to that previously characterized in binocular areas of V1. Covering the left eye assured that all VEPs would originate in the right eye, but since we wanted to compare plasticity in LMC to RBC it was also necessary to prevent visual stimulation of ipsilateral projections from the right eye (Hooks & Chen, 2020). To do this, we translated the stimulus presentation screen right and rotated it laterally until we saw no evoked potentials in the electrode implanted in the RBC (see Fig. 4A and the inset of 4B).

**Figure 4.**
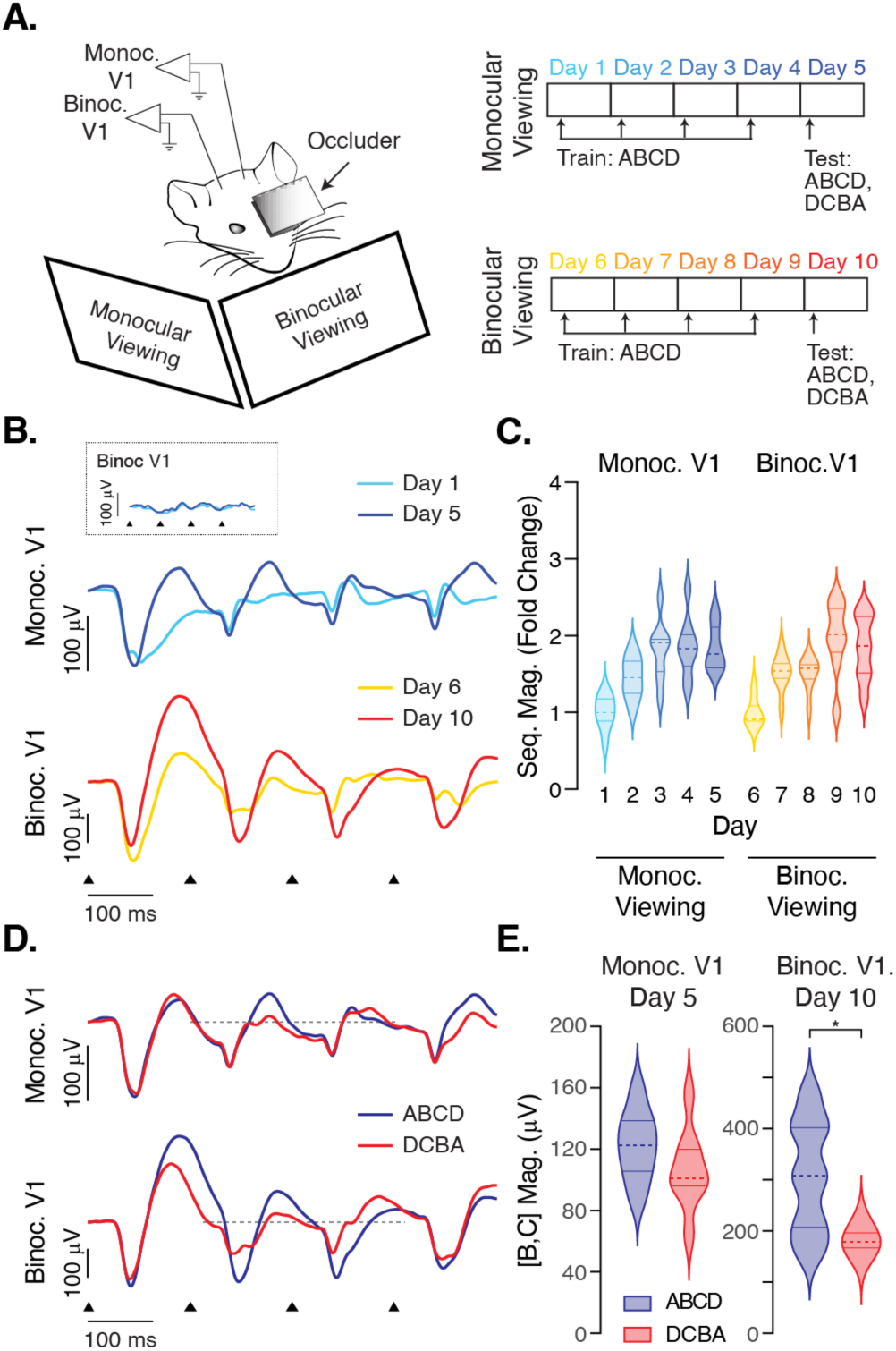
Potentiation in monocular V1 is similar to binocular V1. **A.** Animals (*n* = 8) were implanted with electrodes in monocular V1 in the left hemisphere and binocular V1 in the right. For days 1-5, stimuli were presented only to the monocular cortex (contralateral projections from the right eye were isolated by placing a visual occluder over the left eye and positioning the screen at an angle of approximately 45° off the right side of the snout). For days 6-10 both eyes were open, and the screen was positioned directly in front of the nose. **B.** Monocular stimulation (top) evoked smaller responses than did binocular stimulation (bottom, note scale bar differences) though in both cases the responses potentiated with training. The inset shows LFP recorded in the binocular cortex during monocular training and confirms that evoked activity was restricted to the left monocular cortex. **C.** Violin plots showing evoked responses in monocular (left) and binocular (right) cortex over training normalized by the average group magnitudes on days 1 (monoc) and 6 (binoc). Statistical analysis of fold changes using a mixed model RM-ANOVA reveals that day is highly significant (*F*_4,56_ = 43.058, *p* < 0.001) but that there is no effect of hemisphere (*F*_1,14_ = 0.031, *p* < 0.864) or interaction between the two (*F*_4,56_ = 2.279, *p* = 0.072). These results show that monocular cortex is shaped by visual experience in a manner very similar to binocular cortex, but the level of potentiation does not correlate with M2 expression levels. **D.** The effect of sequence order was tested in both monocular and binocular cortex on days 5 (monocular viewing) and 10 (binocular viewing). Responses to ABCD were larger than DCBA in both hemispheres, though the effect was smaller in monocular cortex **E.** Responses to ABCD were significantly larger than DCBA in binocular cortex (125.9 µV, t(7) = 3.415, *p* = 0.011). ABCD responses were also larger in monocular cortex, but this difference fell just short of significance (15.7 µV, t(7) = 2.239, *p* = 0.060).

Animals were trained with this configuration, with visual responses isolated to the LMC, for four days and tested on day 5. The VEPs recorded in the monocular cortex were similar in shape but smaller in magnitude than those recorded previously in binocular V1, and their magnitude clearly increased with training (Fig. 4B). Starting on day 6, the visual occluder was removed and the screen was repositioned directly in front of the animal so that visual stimulation would activate RBC. Training continued as before on days 6-9 and the animals were tested on day 10. As expected, responses in the binocular cortex potentiated normally (Fig. 4B bottom). To directly compare potentiation in the two hemispheres, we next calculated fold-changes (Fig. 4C) by normalizing quantified sequence magnitudes in monocular recordings to the average response on day 1 and binocular recordings to the average response on day 6 (training day 1 in both cases). We found no statistical difference in potentiation magnitude between monocular and binocular cortex. Average responses to ABCD were larger than DCBA in both hemispheres, though the effect was smaller in monocular cortex (Fig. 4D-E). These findings show that visual experience is sufficient to drive long-term changes in evoked response dynamics in the monocular regions of V1 very similar to those seen in binocular areas (which has not been reported before), with both areas showing similar learning dynamics that saturate at approximately 2-fold of untreated baseline. Whatever functional role increased expression of M2 receptors plays in the monocular cortex, it does not cause experience dependent plasticity to occur faster or with a larger magnitude. This could mean that plasticity requires a baseline of receptor expression and does not increase scale with receptor density beyond this or it might indicate that plasticity is occurring in deeper layers where the M2 expression is approximately equal between monocular and binocular areas.

### Interrupting mAChR signaling after visual stimulation does not prevent sequence potentiation

Sequence potentiation in our experiments is not obvious until hours after stimulus exposure, indicating that some form of consolidation is required to modify V1 circuits. The hippocampus plays a well-known role in memory consolidation, including during sleep, and sequence plasticity does not occur in V1 when the hippocampus is lesioned (Finnie et al., 2021). These findings make it unclear whether mAChR activity is required during visual stimulation or whether relevant signaling continues after stimulation ends. The timing of muscarinic signaling affects episodic memory formation in human subjects who are unable to recall a list of words if scopolamine is delivered before, but not after, presentation of the list (Ghoneim and Mewaldt, 1975; Petersen, 1977). To see whether a similar time dependence holds in V1, we conducted an experiment to determine whether blocking mAChR activity after visual stimulation prevents sequence learning. As before, animals were trained on sequence ABCD for 4 days and tested on day 5 (Fig. 5A). On each training day animals were injected with scopolamine (n=5) or vehicle (n=7) immediately after visual stimulation ended. No drugs were administered on test day. VEPs potentiated normally in both groups with no significant drug effect (Fig. 5B-E). These findings indicate that plasticity requires mAChR activity during stimulus presentation, like human memory experiments.

**Figure 5.**
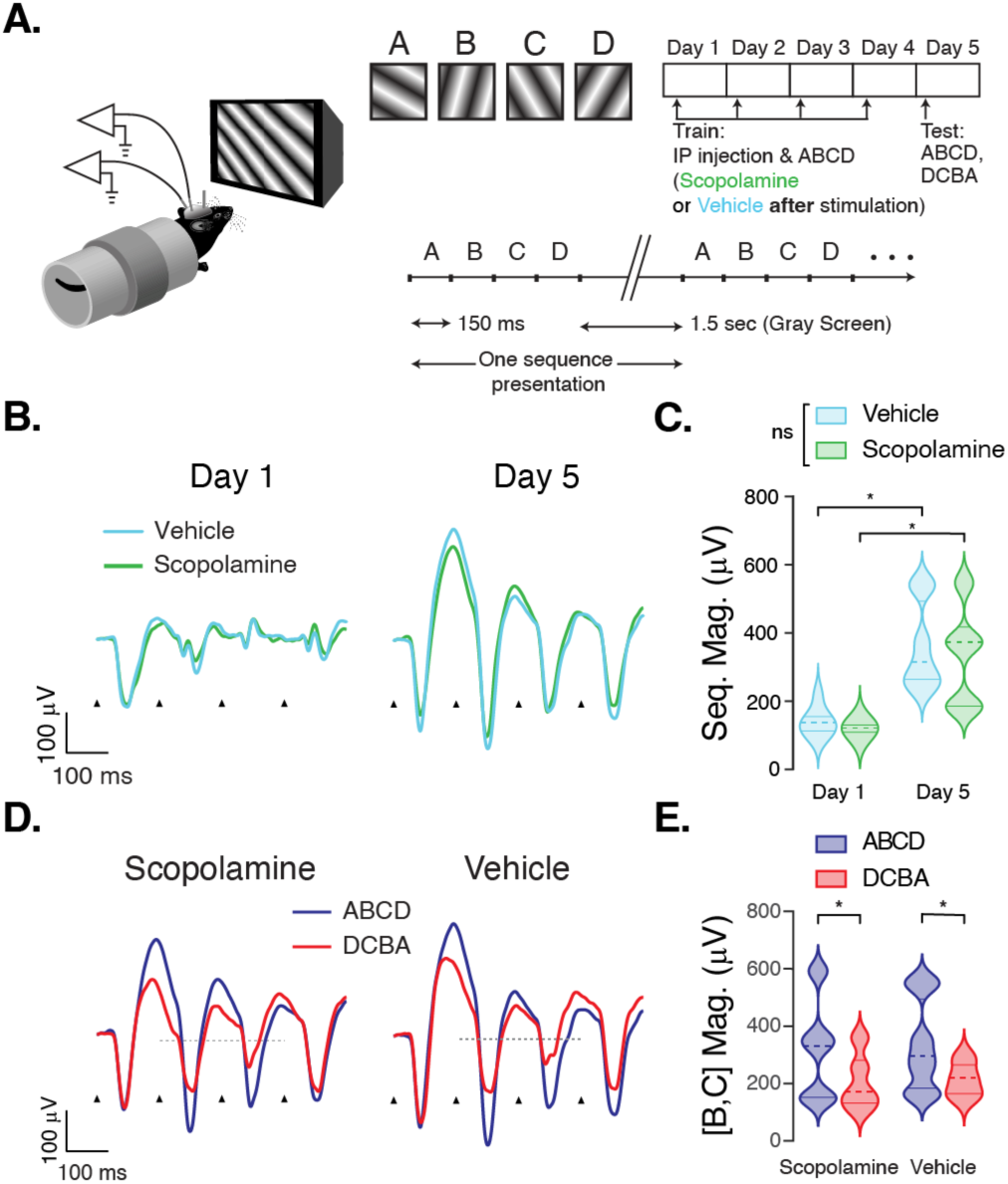
Antagonizing muscarinic receptors after visual stimulation does not prevent spatiotemporal sequence learning in V1. **A.** Mice were treated with either the muscarinic receptor antagonist scopolamine (exp, *n* = 5) or vehicle (ctrl, *n* = 7) immediately after visual stimulation during training. **B.** Sequence-evoked LFPs looked very similar in drug and vehicle-treated animals at baseline (day 1) and after training-induced potentiation (day 5). **C.** Sequence magnitudes recorded on days 1 and 5 showed a significant effect of recording day (*F*_1,9_ = 31.601, *p* < 0.001), but no effect of treatment (*F*_1,9_ = 0.431, *p* =0.528), or significant interaction between treatment and day (*F*_1,9_ = 0.107, *p* =0.751). **D.** Average responses to ABCD (blue) are larger than those to DCBA (red) in drug (left) and vehicle (right) treated mice. **E.** There is a significant effect of element order (*F*_1,18_ = 5.496, *p* = 0.031) but not drug treatment (*F*_1,18_ = 0.114, *p* = 0.740) and no significant interaction between the two (*F*_1,18_ = 0.175, *p* = 0.681). ABCD responses are significantly larger than DCBA in both drug (right-tailed t-test, t(4) = 2.542, *p* = 0.032) and vehicle (t(6) = 2.059, *p* = 0.043) treatment groups.

## Discussion

Cholinergic signaling through mAChRs is implicated in multiple forms of temporal processing in humans and animal models, including spatiotemporal sequence learning, interval-timing (Abner et al., 2001; Balci et al., 2008; Zhang et al., 2019), temporal gap detection and ordinal tasks (Caine et al., 1981; Ison & Bowen, 2000), and auditory discrimination (Seki et al., 2001). Most of these works designed their experiments around scopolamine, which is a non-selective mAChR antagonist. A previous study in rat showed that VEP amplitudes increase following repeated pairings of visual stimulation with electrical stimulation in the basal forebrain, and that this can be prevented with pharmacological antagonists blocking nAChRs, M2 receptors and GABA_A_ receptors (Kang et al., 2015). Our work is the first to isolate the M2 receptor as being required specifically for spatiotemporal plasticity.

M2 receptors can be involved with temporal associative memory tasks via regulation of beta-site APP-cleaving enzyme 1 (BACE1) expression, demonstrated by decreased freezing behavior in trace fear conditioning experiments in mice lacking this enzyme (Ohno et al., 2006; Roßner et al., 2006). M2 knockout mice show performance deficits in different spatial learning tasks such as the Barnes circular maze and T-maze delayed alternation task, including a lack of short-term potentiation and reduced long-term potentiation in the hippocampus after high frequency stimulation that was restored by a GABA_A_ receptor antagonist (Seeger et al., 2004). This, along with a similar role for GABA_A_Rs in Kang et al., 2015 suggests that the M2 receptor’s functional role may involve regulating GABAergic inhibition. Assuming this mechanism holds in V1, it would follow that sequence-evoked potentiation seen in the LFP results in part from dis-inhibition mediated increases in cortical sensitivity rather than increases at excitatory synapses.

Spatiotemporal sequence potentiation looks superficially similar to another form of learning called stimulus selective response potentiation (SRP) that encodes spatial information, though it is mechanistically distinct. SRP requires NMDARs (Cooke and Bear, 2010, 2014) and is not prevented by hippocampal lesion, whereas sequence learning is not affected by NMDAR antagonists (Gavornik and Bear, 2014) and requires an intact hippocampus (see discussion below). Familiar and novel stimuli in the SRP paradigm are characterized by low and high-frequency oscillations in V1 (Hayden et al., 2021), with familiar stimulus evoked PV+ neuron activity decreasing and SST+ activity increasing with learning. Given that SRP requires NMDARs in PV cells for plasticity to occur (Kaplan et al., 2016), it is not clear whether to expect a similar finding in sequence learning. It should also be noted that behavioral state, including locomotion, exerts a large effect on cortical processing (Fu et al., 2014; Stringer et al., 2019; Heintz et al., 2022) and can modify various forms of stimulus selective plasticity (Montgomery et al., 2022). It seems likely that similar effects will also hold for spatiotemporal plasticity, though many details remain to be explored.

Our identification of somatic M2 expression in SST+ neurons suggests that these cells may play a role in this process. One possibility is that M2 receptor mediated plasticity modulates SST neurons to implement a “temporal surround suppression,” akin to their role in spatial suppression (Niell and Scanziani, 2021), that biases responses towards learned spatiotemporal patterns. It is not clear what this would look like in practice, or whether it would even be possible to see by measuring somatic spikes. The natural assumption might be that M2-related plasticity would cause SST neurons to fire less, but since M2 receptors are primarily presynaptic autoinhibition could decrease neurotransmitter release at individual synapses based on localized ACh levels without effecting overall firing rates. It should also be noted that other cell types could also play an important role in mediating plasticity. For example, M2AChR expression has been localized to PV+ cells in the rat entorhinal cortex using RNA amplification following single cell dissection (Chaudhuri et al., 2005) and there is no reason to expect that all the M2 receptors in the neuropil are located on neurites from SST+ cells. M2 receptors can play a variety of functional roles including hyperpolarizing the neuronal membrane by opening inward rectifying K^+^ channels (Brown, 2010) and preventing the neurotransmitter release at GABAergic terminals (Salgado et al., 2007). Cholinergic activity can enhance thalamocortical inputs, improve attention, etc. (Sarter and Parikh, 2005; Sarter et al., 2005; Groleau et al., 2015) and any of these processes could impact plasticity and be modified by M2 receptor action. Given the relation between mAChR mediated plasticity and GABA receptors found in other studies, it will be important to identify the exact role M2 plays in V1 to clarify its role in shaping temporally specific plasticity.

One interesting question is how scopolamine-induced amnesia (Caine et al., 1981) overlaps with hippocampal-dependent memory formation. It is well established that the hippocampus plays a critical role in the creation of long-term episodic memories, and damage to the medial temporal lobe can prevent sequence potentiation in V1 (Finnie et al., 2021). Since hippocampal activity during sleep is involved with memory consolidation mechanisms (Reyes-Resina et al., 2021), these results could indicate that V1 consolidation explicitly requires an intact hippocampus. Our findings, however, suggest that the muscarinic signaling necessary to initiate plasticity occurs during visual stimulation itself and call into question whether there is an active role for the hippocampus outside of this time window. An alternative possibility is that hippocampal projections to the medial septum via the fornix are required for normal cholinergic activity in the basal forebrain and that sequence activity is indirectly blocked when hippocampal lesions interrupt this activity. Given the recently discovered modulation of V1 dynamics by hippocampal place representations (Saleem et al., 2018; Niell and Scanziani, 2021), it will be important to determine the extent to which plasticity in local cortical circuits are directly shaped by activity in the hippocampus, indirectly by hippocampal-dependent control of neuromodulators, or require hippocampal activity during sleep (Klinzing et al., 2019).

Other studies have also found M2 receptors expressed in interneurons (Volpicelli and Levey, 2004; Groleau et al., 2015). M2 receptors specifically have been found in SST positive neurons in the rat amygdala, rat hippocampus and rat auditory cortex (Hajos et al., 1997; McDonald & Mascagni, 2011; Salgado et al., 2007), and non-specific colocalization of mAChRs with SST has also been reported previously in mouse V1 (Chen et al., 2015). Interestingly, mAChRs are involved in mismatch negativity (MMN) in the auditory system (Pekkonen et al., 2001; Schöbi et al., 2021; Weber et al., 2021) and SST neurons have been implicated in visual perception and mismatch detection (Song et al., 2020; Hamm and Yuste, 2016; Attinger et al., 2017). MMN is an electrophysiological response generated in the brain when an expected stimulus is replaced by a deviant or oddball stimulus, and while the timeframe of this effect is quite different from the multi-day plasticity experiments used here, there may be a relation in how cholinergic modulation and inhibitory circuits recognize novelty. Reduced MMN during the processing of sensory stimuli is a hallmark of schizophrenia (Baldeweg et al., 2006; Inami et al., 2008; Garrido et al., 2009; Inami and Kirino, 2019) and reduction of SST mRNA and abnormal SST neuron localization has also been observed in people with schizophrenia (Morris et al., 2008; Urban-Ciecko and Barth, 2016). SST neurons form synapses on both Vasoactive Intestinal Peptide (VIP) interneurons and pyramidal excitatory neurons, and a recent study focusing on the role of visuomotor experience in shaping predictive coding in V1 found specific roles for both SST and VIP cells in signaling expectation violations (Attinger et al., 2017). Cumulatively, these studies highlight the role of muscarinic signaling in temporal predictive processes, may indicate a direct link between mAChR signaling in SST neurons and MMN suppression in schizophrenia, and possibly represent a mechanistic locus that could be probed in V1 circuits.

## Acknowledgements

We thank all the mice that gave their lives for this research, without their sacrifice this paper wouldn’t have existed. We thank Gianna Ferron of the Gavornik Lab for her support running experiments, and Todd Blute of the Boston University Biology Imaging Core for help using the fluorescence microscope. Finally, thanks to the great folks at Microsoft who after decades of tireless work still don’t know how to make a word processing program where it is possible to cut/paste without destroying all the formatting. This work was supported by grants from NIMH R00MH099654 and NEI R01EY030200.

## Author Contributions

Conception and experimental design: S.S. and J.P.G. Surgeries, experiments, and data collection: S.S. and C.M.J. Data analysis: S.S., C.R-M. and J.P.G. Writing and figures: S.S., C.R-M., C.M.J, and J.P.G.

## Notes

https://gavorniklab.bu.edu/supplemental-materials.html

### Competing Interest Statement

The authors have declared no competing interest.

### Summary of Updates

Additional experiments to provide a positive control for M1 antagonist. Figure revisions to include supplemental in main body.

